## Supplementary Figures for "Nonlinear feedback modulation contributes to the optimization of flexible decision-making"

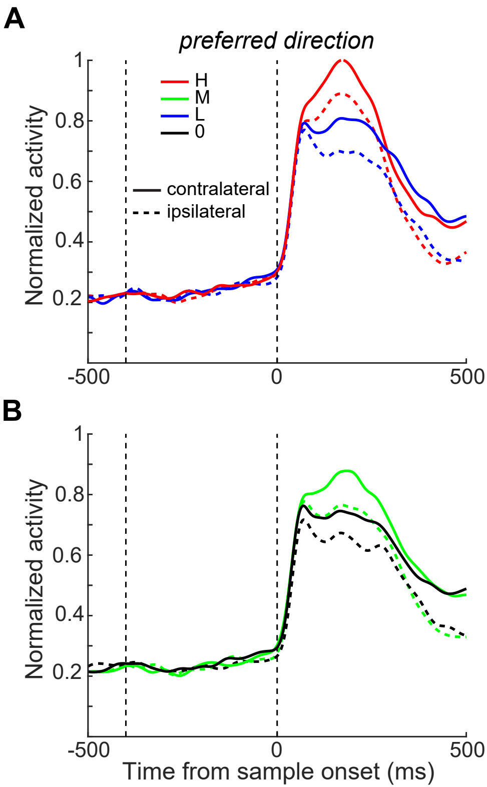


**Figure S1**. The comparisons of LIP activity between the CT and IT conditions. The averaged population activity responded to motion stimuli in both CT and IT conditions are shown separately for different motion coherence levels. The averaged LIP activity to the preferred (**A-B**) and nonpreferred (**C-D**) motion directions were shown separately. Different colors denote different motion coherence levels. The two vertica dashed lines denote the time of target and motion stimulus onset, respectively.

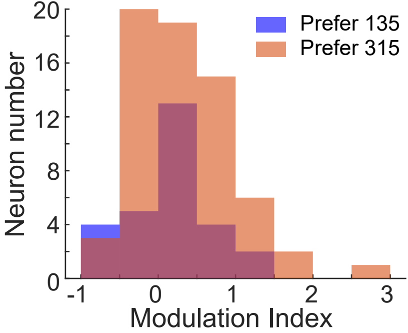


**Figure S2** There was no systematic relationship between direction preference and saccade-related modulation in LIP neurons' responses to motion stimuli. A modulation index was computed for each neuron to quantify differences in motion direction selectivity between the CT and IT conditions. The distribution of modulation indices was then compared between neurons preferring 315° motion direction (red) and those preferring 135° motion direction (blue).

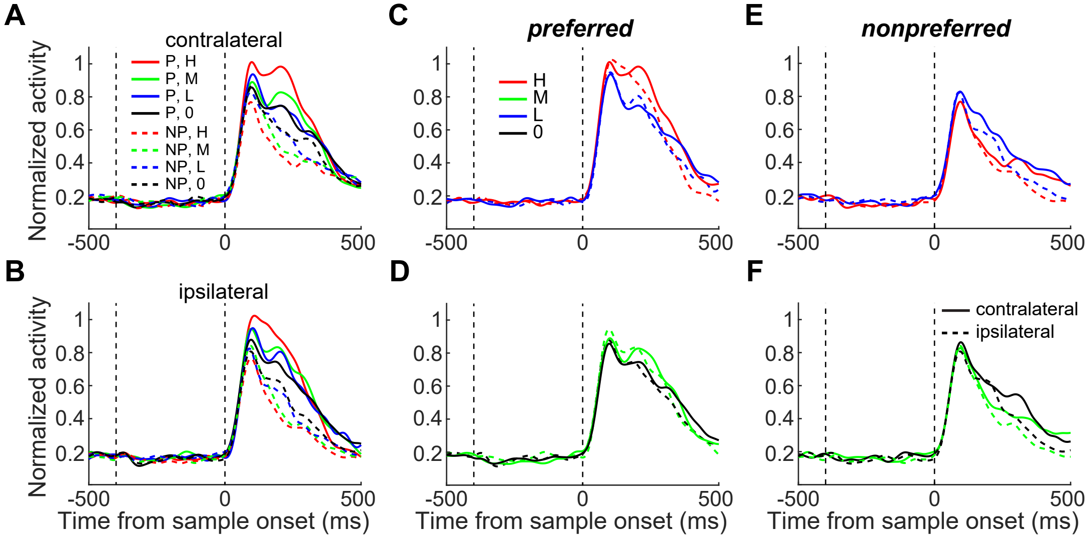


**Figure S3.** The comparison of LIP activity between the CT and IT conditions. This figure only includes the data sessions in which the saccade targets were aligned close to the vertical direction. (**A-B**) The averaged population activities in both the CT (**A**) and IT (**B**) conditions are shown separately for each motion direction and coherence level. (**C-D**) The comparisons of LIP activity responded to the preferred motion direction between the CT and IT conditions are shown for different coherence levels. (**E-F**). The comparisons of LIP activity responded to the nonpreferred motion direction between the CT and IT conditions.

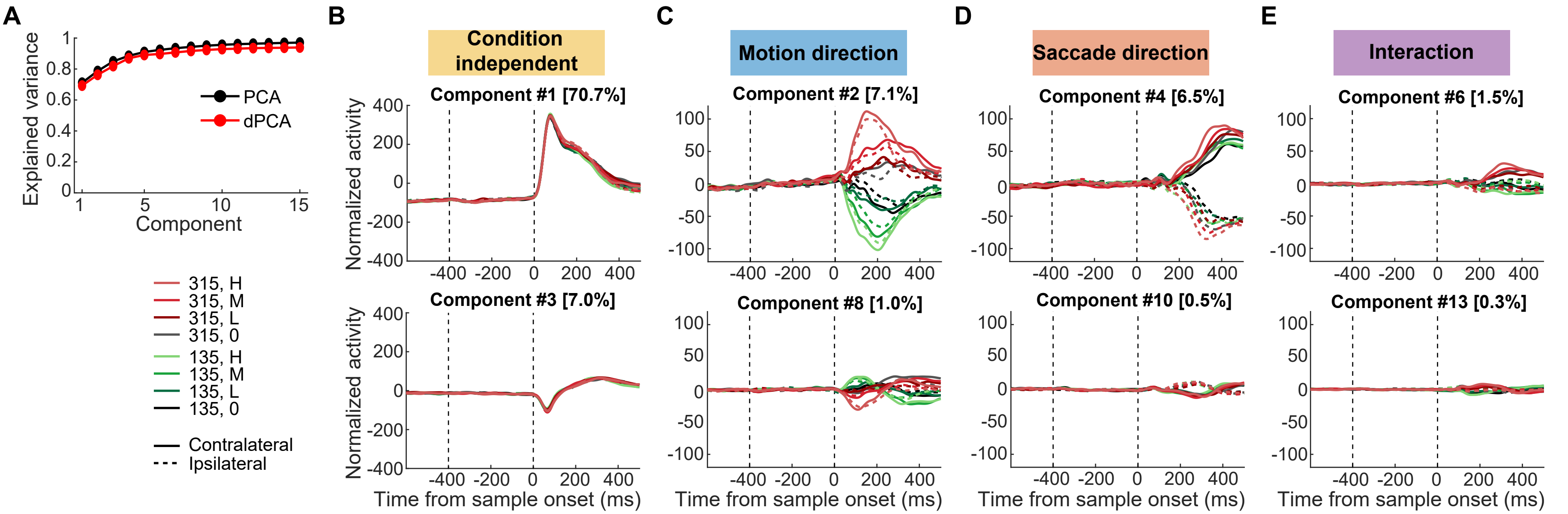


**Figure S4**. The motion and saccade representations in LIP shown by dPCA analysis. LIP population activity was decomposed into four task-related variables: motion direction, saccade direction, motion-saccade interaction, and timing (condition-independent). (**A**) Cumulative variance explained by PCA (black) and dPCA (red) for LIP population activity responded to motion stimuli. Only the first 15 principal components (PCs) were shown. dPCA explains almost the similar amount of variance as standard PCA . (**B-E**) Demixed principal components. The upper row: the first demixed PCs of LIP population activity corresponding to the four variables. The lower row: the second demixed PCs of LIP population activity. In each subplot, LIP population activity is projected onto the respective dPCA decoder axis, so that there are 16 lines corresponding to 16 conditions (2 motion directions × 4 coherence levels × 2 saccade directions). Different colors represent different motion directions. Solid and dashed lines represent CT condition and IT condition, respectively. The two vertical dashed line mark the time of target and motion stimulus onset, respectively.

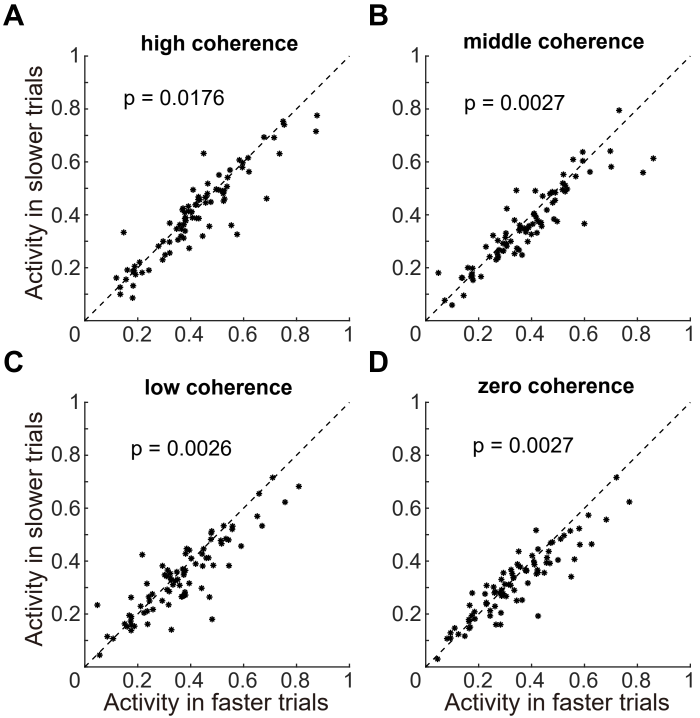


**Figure S5.** The comparison of LIP activity to the preferred motion direction between the faster and slower RT trials in the CT condition. Data from each coherence level is shown separately. Each dot denotes the averaged activity of a single neuron.

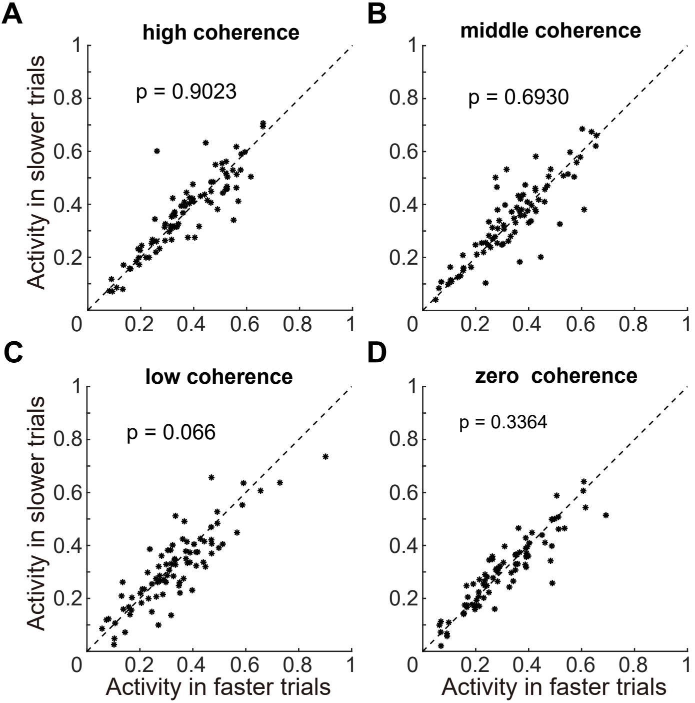


**Figure S6.** The comparisons of LIP activity to the preferred motion direction between the faster and slower RT trials in the IT condition, which are shown in the same format as in Figure S4.

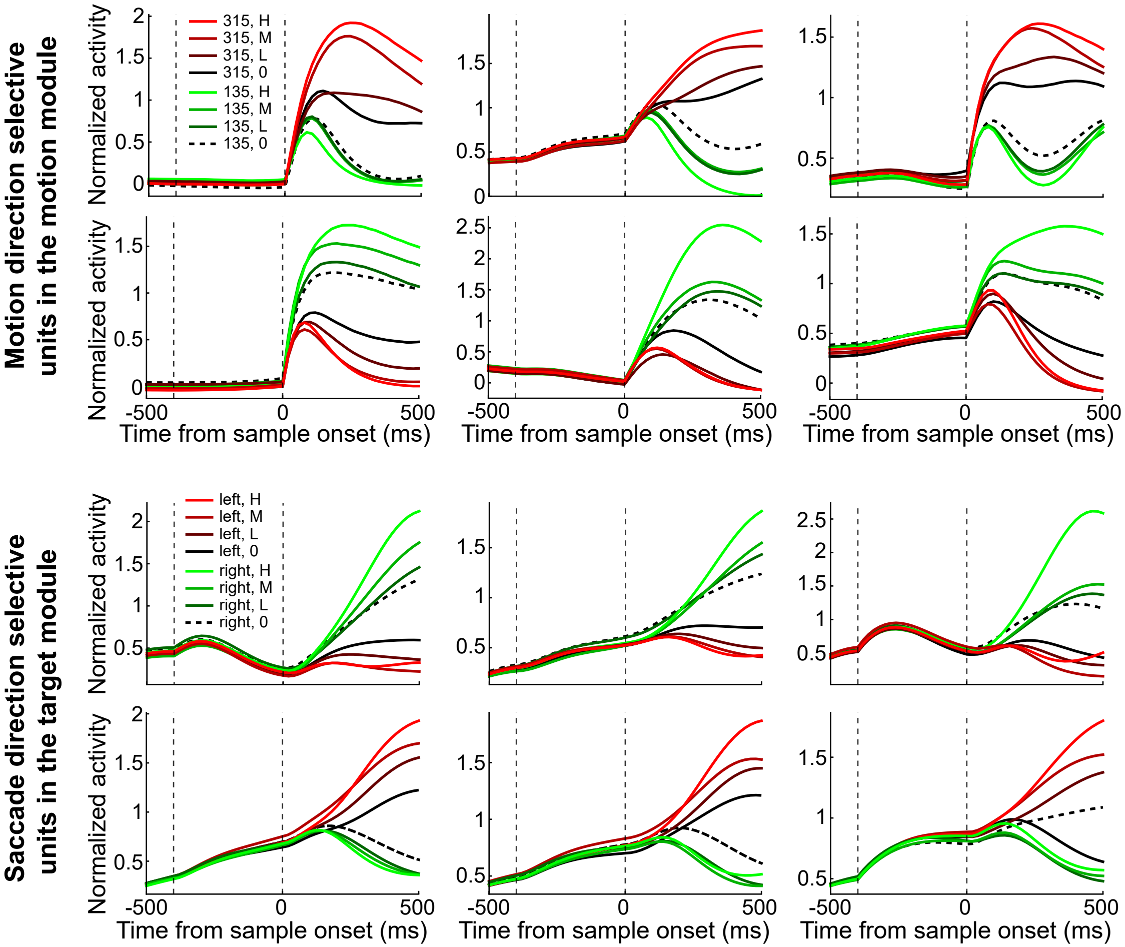


**Figure S7.** Examples of unit activity in the example network. The two upper rows show the activities of 6 example units in the motion module. Different colors denote different motion directions, and different shades denote different coherence levels. The zero coherence trials were grouped based on the network's choices. The two lower rows show the activities of 6 example units in the target module. Different colors denote different saccade directions.

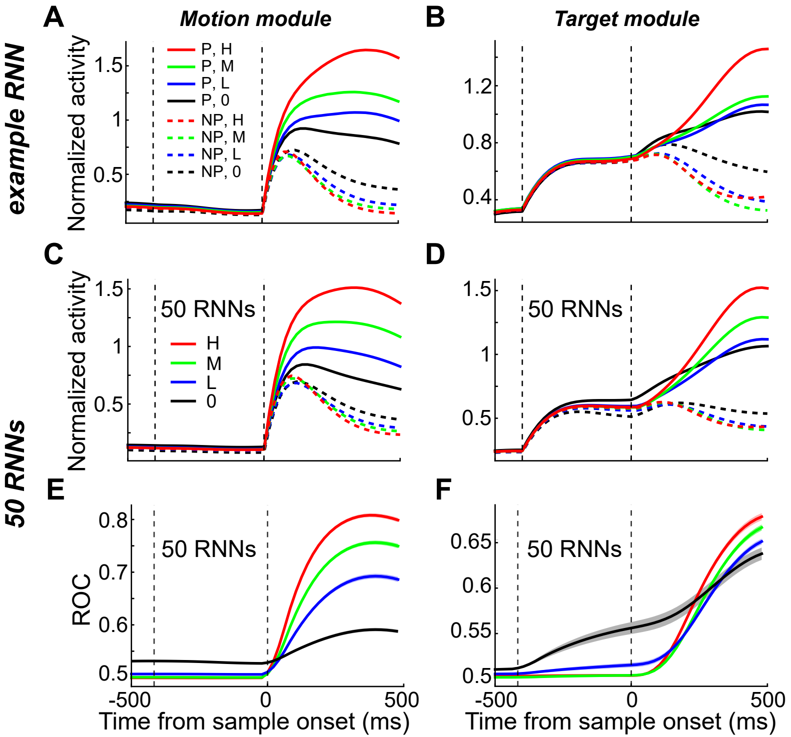


**Figure S8**. Population level unit activity across all trained RNNs. (**A**) The averaged population activity of all motion direction selective units in the motion module of the example RNN. (**B**) The averaged population activity of units in the target module of the same example RNN is shown for different saccade directions and motion coherence levels. (**C-D**) The averaged population activity of all 50 trained RNNs is shown in the same format as in a-b. (**E-F)** An ROC analysis was used for quantifying the motion DS for the motion module (h) and saccade DS in the target module for all 50 trained RNNs.

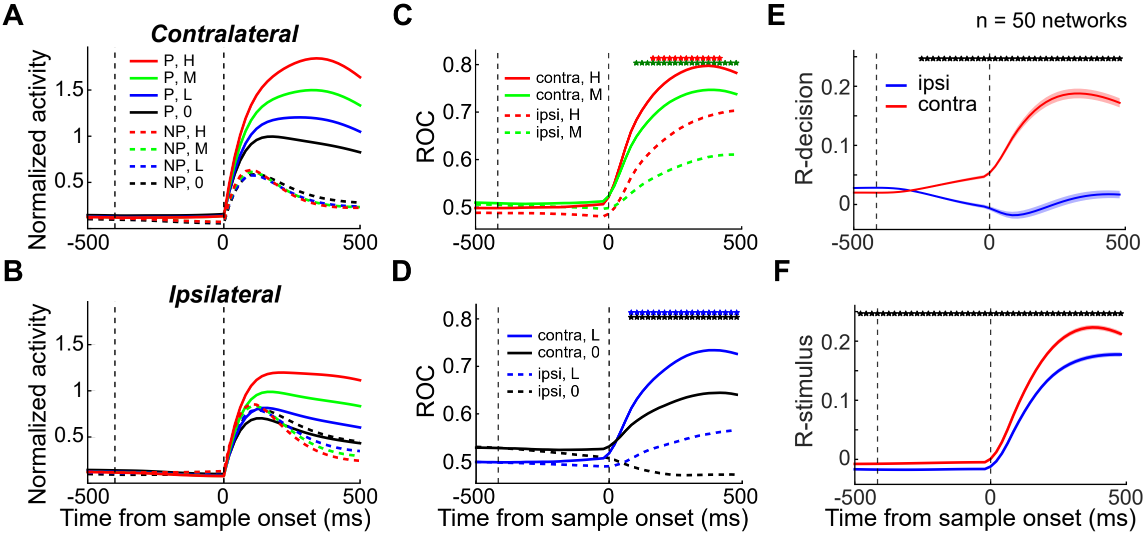


**Figure S9.** The motion direction selectivity in the motion module was significantly modulated by later saccade choice in the RNNs. Only the units in the motion module were included, and data from all the trained RNNs was pulled together. (**A-B**) The averaged population activities of all direction-selective units in all the trained RNNs are shown separately for CT condition (**A**) and IT conditions (**B**). (**C-D**) The motion DS in the motion module of all the trained RNNs was quantified by ROC analysis for all 4 motion coherence levels. Solid and dashed lines denote data in the CT and IT conditions, respectively. The color dots denote the time points for which there was significant difference between CT and IT conditions ($p < 0.01$, paired t-test). (**E-F**) Partial correlation analysis. The averaged value of r-decision (**E**) and r-stimulus (**F**) across all the trained RNNs are compared between IT and CT conditions.

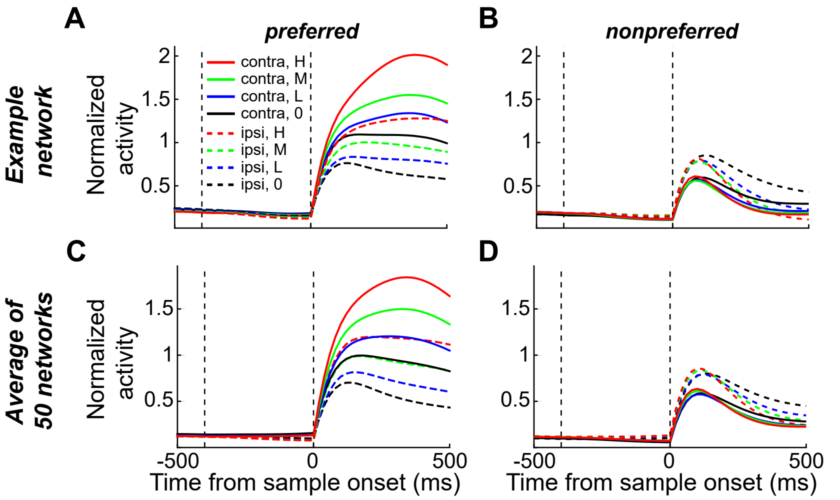


**Figure S10**. The comparison of unit activity in the motion module of the RNNs between the CT and IT conditions. (**A-B**) The averaged population activities of the example RNNs responded to the preferred (**A**) and nonpreferred (**B**) motion directions are shown separately for the CT (solid) and IT (dashed) conditions. (**C-D**) The comparisons of unit activity averaged across all the trained RNNs between the CT and IT conditions are shown for different coherence levels.

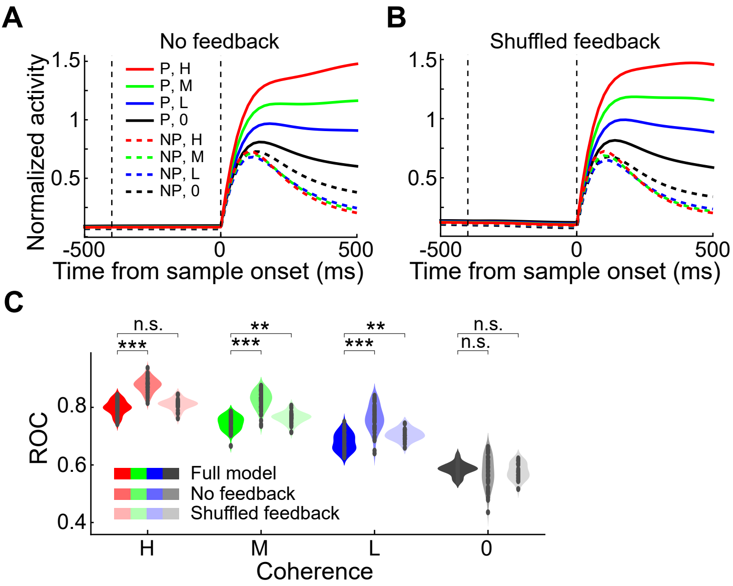


**Figure S11**. The motion DS in the motion module of the trained RNNs did not decrease after inactivating feedback connectivity. (**A**) The averaged population activity of units in the motion module of all RNNs after ablating feedback connections were shown separately for different motion directions and coherence. (**B**) The averaged population activity of units in the motion module of all RNNs after disrupting feedback connections were shown in the same format as in a. (**C**) The averaged motion DS in motion module of the full-model RNNs, RNNs without feedback connections and RNNs with disrupted feedback connections were compared separately for different motion coherence. An ROC analysis was used for quantifying the motion DS. (Paired t-test: **, P < 0.01; ***, P < 0.001; n.s., not significant).

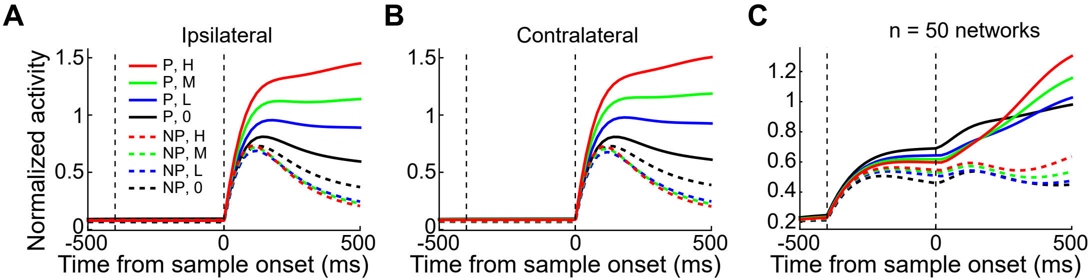


**Figure S12**. The averaged population activity of all trained RNNs without feedback connections. (**A-B**) The averaged population activity of all direction-selective units in the motion module of all the trained RNNs after ablating the feedback connections is shown for different motion directions and coherence levels. Data form both CT condition (**A**) and IT conditions (**B**) were shown separately. (**C**) The averaged population activity of units in the target module of all the trained RNNs after ablating feedback connections is shown for different saccade directions and motion coherence levels.

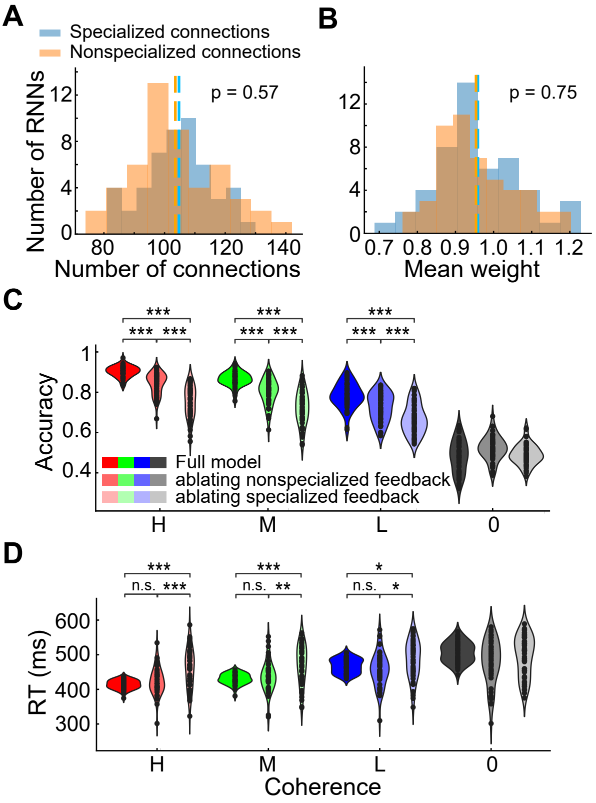


**Figure S13** The effects of pattern-specific ablation of the feedback connections on the RNNs' behavior performance. The feedback connections in either the specialized group or nonspecialized group were ablated separately when tested with the untrained motion stimuli. (**A**) The comparison of the numbers of feedback connections between selectivity-specialized and nonspecialized groups. The distributions of the total number of feedback connections for all 50 RNNs were shown separately for specialized and nonspecialized groups. The blue and brown vertical dashed lines denote the mean values across all 50 RNNs for specialized connections and nonspecialized connections, respectively. (**B)** The comparison of the mean weights of feedback connections between selectivity-specialized and nonspecialized groups. (**C**) The comparison of the networks' performance accuracy. The performance accuracies of the full-modell RNNs, RNNs without selectivity-specialized feedback connections and RNNs without nonspecialized feedback connections were shown separately for different motion coherence. (**D**) The comparison of the networks' RTs. The RTs of the full-modell RNNs, RNNs without selectivity-specialized feedback connections and RNNs without nonspecialized feedback connections were shown separately. (Paired t-test: *, P < 0.05; **, P < 0.01; ***, P < 0.001; n.s., not significant).

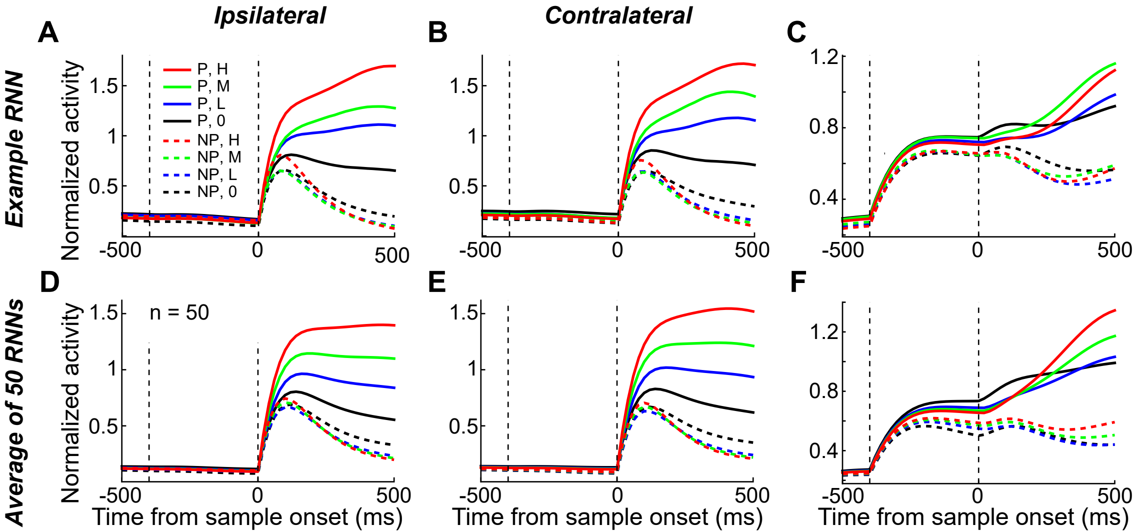


**Figure S14**. The averaged activity of RNNs with disrupted feedback connectivity. (**A-B**) The averaged activity of units in the motion module of the example RNN after disrupting the feedback connectivity is shown for different motion directions and coherence levels. Data in both IT (**A**) and CT (**B**) conditions were shown separately. (**C**) The averaged activity of units in the target module of the example RNN after disrupting the feedback connectivity is shown separately for different saccade directions and motion coherence levels. (**D-E**) The averaged population activity of all direction-selective units in the motion module of all the trained RNNs after disrupting the feedback connectivity is shown separately for both IT condition(**D**) and CT conditions (**E**). **(F**) The averaged population activity of units in the target module of all the trained RNNs after disrupting feedback connectivity is shown for different saccade directions and motion coherence levels.

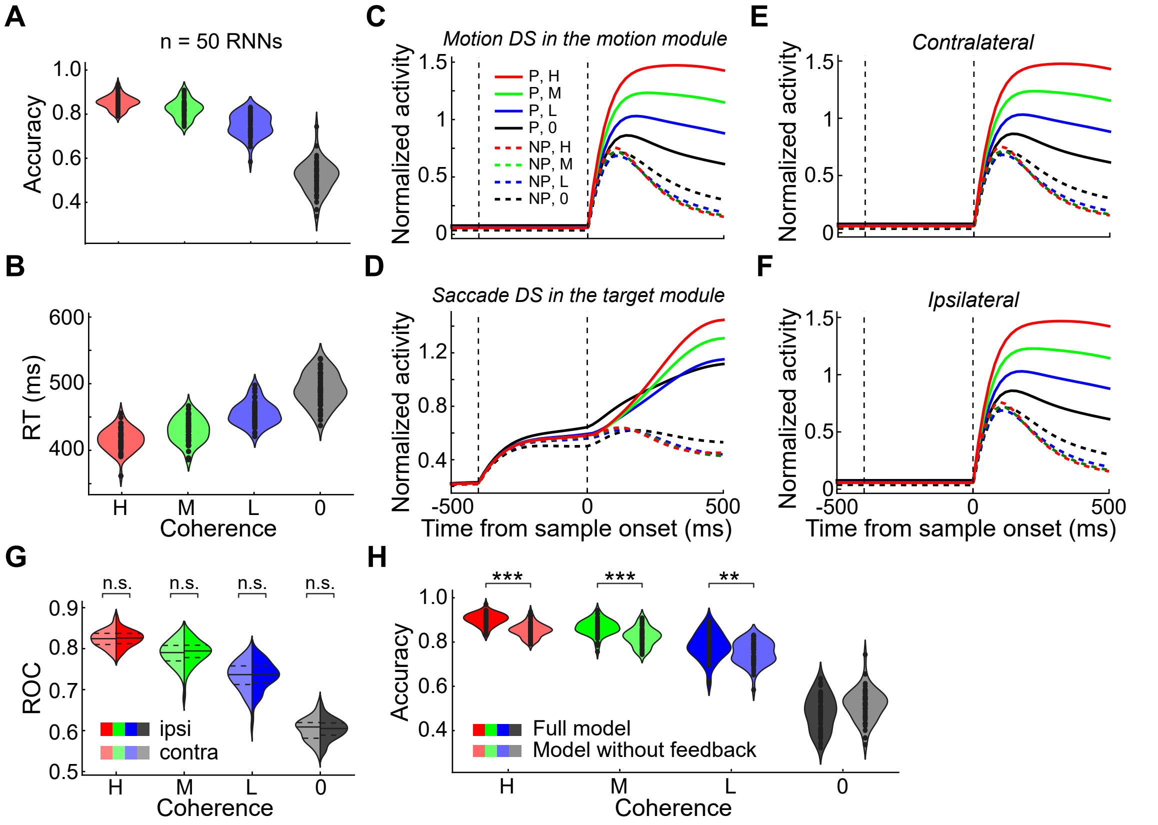


**Figure S15**. The behavioral performance and unit activity in the RNNs that were initialized without feedback connectivity. (**A-B**) The performance accuracies (**A**) and reaction times (**B**) of all 50 trained RNNs are shown separately for each motion coherence levels. (**C**) The averaged population activity of all motion direction selective units in the motion module is shown for each motion direction and coherence level. Data from all 50 networks were averaged. (**D**) The averaged population activity of all saccade direction selective units in the target module is shown for each saccade direction and coherence level. (**E-F**) The averaged population activities of all direction-selective units in the motion module of all 50 RNNs were shown separately for CT (**E**) and IT conditions (**F**). (**G**) The comparisons of the motion DS between the CT and IT conditions (quantified by ROC analysis) in the motion module of all 50 RNNs are shown separately for different coherence levels. (**H**) The comparisons of the performance accuracies between full-model RNNs and RNNs trained without feedback connections (Paired t-test: **, P < 0.01; ***, P < 0.001; n.s., not significant)..

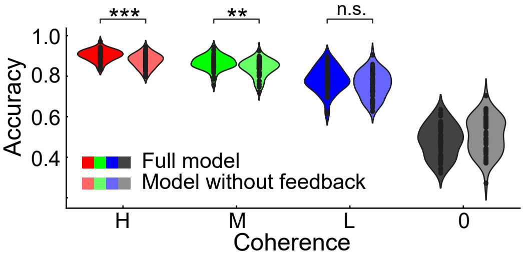


**Figure S16**. The performance accuracies of the additional RNNs trained without feedback connectivity are shown separately for each motion coherence level. These RNNs were initialized with greater recurrent connection probabilities than the full-model RNNs, so that the number of the total trainable connection weights matched that in the full-model RNNs. (Paired t-test: **, $P < 0.01$; ***, $P < 0.001$; n.s., not significant).
